# Anti-malarial contact dependent blocking of transmission of *Plasmodium vivax* by *Anopheles darlingi* mosquito vector

**DOI:** 10.1101/2025.09.11.675498

**Authors:** Jéssica E. A. Kassupá, Alice O. Andrade, Alessandra S. Bastos, Gabriel L. L. Moura, Marina L. Rocha, Nascimento Martinez Leandro do Na, Elisandra Kathellen da Silva Afonso, Wallyson de Jesus da Costa, Daniel Y. Bargieri, Carolina Bioni Garcia Teles, Jansen F. Medeiros, Nirbhay Kumar, Ana Carolina Ramos Guimarães, Douglas G. Paton, Flaminia Catteruccia, Maisa S. Araujo

## Abstract

Malaria, caused by protozoa of the genus *Plasmodium* and transmitted to humans through the bite of mosquitoes of the genus *Anopheles*, remains a public health problem. Long-Lasting Insecticide -treated bed Nets (LLINS) and Indoor Residual Spraying (IRS) represent the main vector control measures for malaria prevention. However, to address the concerns of mosquito resistance to pyrethroids, other malaria control strategies are being explored for effectively blocking malaria transmission by eliminating or reducing the parasite in the vector. This study evaluated the use of antimalarials through tarsal contact of female *Anopheles darlingi* infected with *Plasmodium vivax* via a Direct Membrane Feeding Assay (DMFA). Female *An. darlingi* were exposed tarsally using Petri dishes impregnated with antimalarials at 1 mmol/m^2^ for exposure times of 6 or 60 minutes. Among the antimalarials evaluated were Atovaquone (ATQ), Tafenoquine (TQ), Chloroquine (CQ), Mefloquine (MQ), Primaquine (PQ), and the compound Nanchangmycin (NCG). Atovaquone was the only antimalarial evaluated before and after DMFA at exposure times of 60 min and 6 min. The results demonstrate complete elimination of *P. vivax* in female *An. darlingi* exposed to ATQ by tarsal contact 60 min before infection. ATQ was also effective 6 min before or after infection, reducing infection prevalence. In addition, MQ also significantly reduced infection intensity, but there was no difference in infection prevalence. No significant differences were observed for the other antimalarials.

**Author Summary:** Malaria caused by *Plasmodium viva*x is the most prevalent in the Amazon region, with *Anopheles darlingi* as its main vector. Mosquito resistance to pyrethroid insecticides, already described in African countries, also raises an alarm for areas endemic for vivax malaria. Given this scenario, strategies that involve blocking parasite transmission have proven effective. Our study involved the transmission-blocking potential of antimalarials administered via tarsal contact to *An. darlingi* infected with *P. vivax*. Tarsal exposure involves the direct contact of the mosquitoes’ tarsi with surfaces impregnated with antimalarials. We speculate that in a real-world setting, this approach could be translated by treating surfaces like bed nets, eaves, or resting sites with the compounds, exploiting the natural resting and host-seeking behaviors of mosquitoes that bring their tarsi into contact with these treated substrates. If the drugs/compounds penetrate the cuticle, they could impact the parasite’s biological cycle within the vector. Our results confirmed this potential, Atovaquone was able to eliminate or reduce *P. vivax* from the midguts of *An. darlingi* at different exposure times, demonstrating successful uptake through the tarsi. Mefloquine was also reduced parasite intensity via tarsal contact. These findings reinforce the potential of this approach as a complementary tool in malaria control.

## Introduction

Malaria is a disease caused by *Plasmodium* parasites transmitted to humans through bites of infected *Anopheles* mosquitoes. In 2023, malaria accounted for 263 million cases and 597,000 deaths worldwide, representing a persistent public health challenge [1]. The Global Technical Strategy for Malaria (GTS) through 2030 outlines malaria elimination measures targeting both parasites and vectors. These efforts seek to interrupt local transmission by reducing the human parasite reservoir and addressing outdoor transmission including chemotherapeutic interventions to block the malaria transmission cycle. In this context, research is expected to lead to new interventions such as vaccines, new and more effectives drugs and combinations, novel insecticides or combinations, repellents, toxic baits for vectors and other innovations in vector control [2].

Vector control, primarily through Indoor Residual Spraying (IRS) and Long-Lasting Insecticide -treated bed Nets (LLINs), is an important malaria prevent strategy. Until recently, LLINs relied solely on a single class of insecticide, pyrethroids [1,3]. However, emergence of pyrethroids resistance has presented a primary threat to the long-term viability of LLINs [4,5], driven by its rapid geographic spread [2,4]. This highlights the need for new strategies targeting to control malaria transmission with novel active approaches [6–10] to address pyrethroid resistance and enhance the effectiveness of current malaria control strategies [8,9]. To mitigate the resistance issue includes two new classes of dual active-ingredient: pyrethroid-clorfenapyr, which combine pyrethroid and pyrrole insecticide to enhance the net’s lethality [11], and pyrethroid-pyriproxyfen nets, which pair a pyrethroid with an insect growth regulator (IGR) [12] have been developed for impregnating LLINs. Both combinations aim to improve efficacy against pyrethroid-resistant mosquitoes [8]. To contribute to these strategies, Paton et al. [13] generated a new strategy that at least partially overcomes the challenge of insecticide resistance in LLINs by blocking parasite transmission by the *Anopheles* mosquito. Their study exposed *An. gambiae* (s.s.), a primary malaria vector in Africa, to the antimalarial atovaquone (ATQ) prior to infection with *P. falciparum*.

This exposure via tarsal contact eliminated parasites from the mosquitoes’ midguts and reduced both the intensity and prevalence of *P. falciparum* infection in pyrethroid-resistant *An. coluzzi* [14]. In a recent study, they screened additional compounds with the ability to block infections, identifying a compound combination that retained full anti-plasmodial activity even after incorporation into bed net-like substrates. Overall, these studies validate this approach as a promising malaria control tool [15].

The strategy of exposing *Anopheles* females to antimalarials before and also after *Plasmodium* infection [13–15] is based on their tendency to feed at night, when people sleep under mosquito nets. Moreover, after feeding, females rest on internal walls, likely to regain flight capacity and/or digest the blood meal before reaching a gravid state [16]. This feeding and resting behavior is characteristic of certain *Anopheles* species that exhibit more endophagic and endophilic [17] as well as anthropophilic traits, such as *An. darlingi*, a primary malaria vector in the Amazon region [18,19]. Reorienting the use of LLINs and IRS to deliver antimalarial through tarsal contact addresses key challenges associated with drug resistance in parasites and insecticide resistance in mosquitoes [14,20]. Additionally, this approach offers novel opportunities for vector targeted drug delivery [9], disrupting sporogonic development, eliminating the parasites within the mosquito, and thereby blocking transmission to humans. In this context, the present study investigates this transmission-blocking strategy by evaluating the impact of antimalarials and other compounds using the *P. vivax-An. darlingi* model through tarsal exposure, a malaria species that is comparatively neglected, harder to study due to challenges in continuous *in vitro* culture, and likely to be more difficult to eliminate than *P. falciparum*.

## Results

### Exposure to Atovaquone substantially reduces infection of *Anopheles darlingi* with *Plasmodium vivax* isolates

To test whether ATQ, a parasite cytochrome-b inhibitor, could inhibit *P. vivax* development in mosquitoes, we allowed *An. darlingi* females mosquitoes to rest on a glass substrate coated with ATQ immediately before *P. vivax* infection via DMFA. Exposing *An. darlingi* to ATQ at 1 mmol/m^2^ for 60 min resulted in 100% inhibition of *P. vivax* oocysts development after the infectious blood meal, whereas control mock-exposed mosquitoes exhibited a high prevalence and intensity of infection (Fig 1A). In a subsequent experiment, the exposure time of mosquitoes to ATQ was reduced to 6 min. Although this did not completely block transmission, both prevalence and intensity of infection were significantly reduced (See Fig 1B). The transmission reduction activity (TRA) was 97.49%, while the transmission blocking activity (TBA) was above 73.97% (S1A Table).

**Fig 1.**
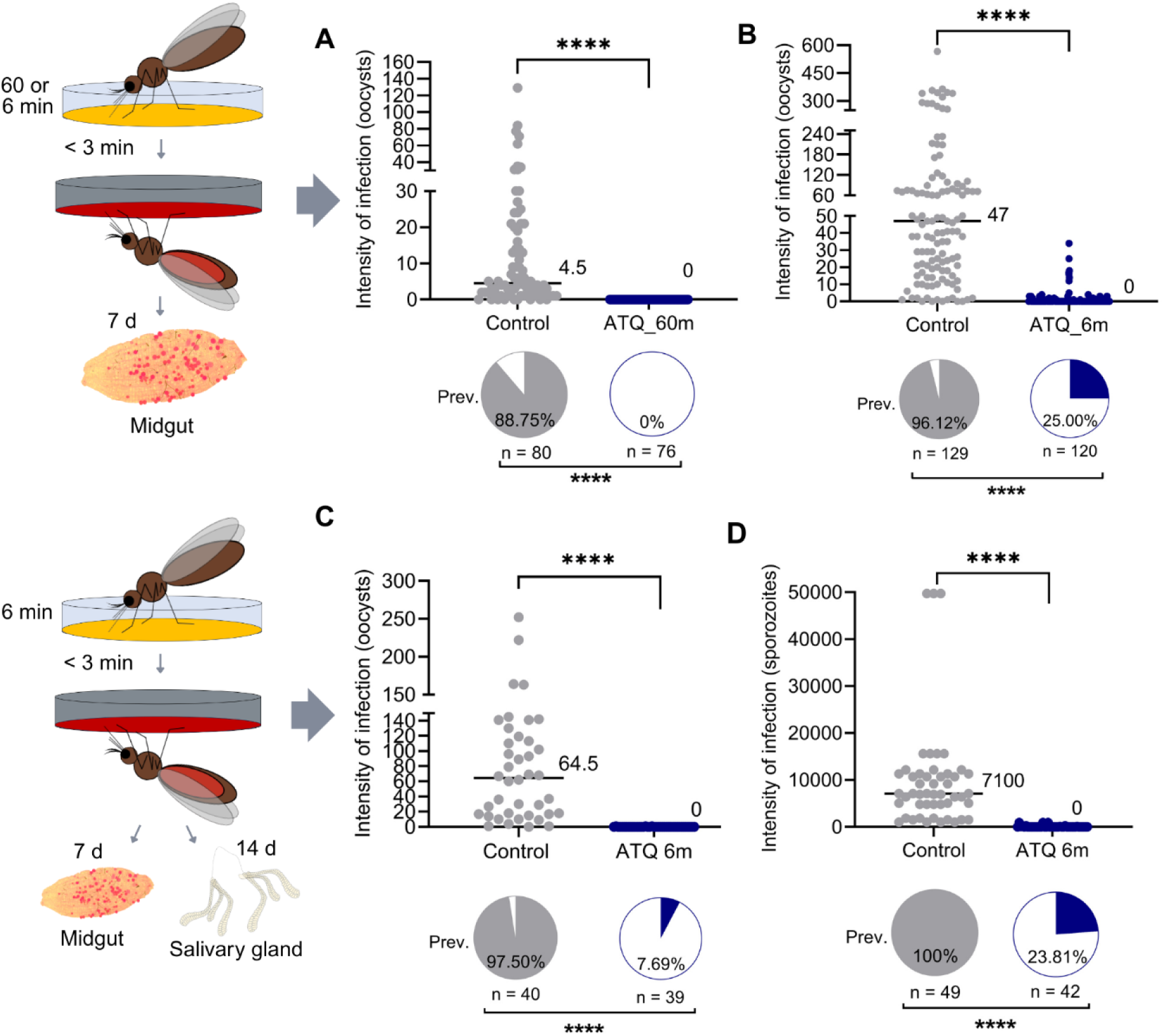
*Anopheles darlingi* exposure to Atovaquone (ATQ) affects *Plasmodium vivax* development. A) *Plasmodium vivax* parasite development was blocked (0 oocyst intensity and 0% prevalence of infection; shown in the pie charts in female mosquitoes exposed to ATQ at 1 mmol per m^2^ for 60 min immediately before infection. Prevalence (Prev.): two-sided chi-squared test, *n =* 156, degrees of freedom (df) = 1, χ2 = 123.8, \*\*\*\**P* < 0.0001. Intensity: two-sided Mann-Whitney *U* test, *n* = 156, df = 1, *U =* 342, \*\*\*\**P* < 0.0001. The exposure method is shown in the graphic: orange represent ATQ coated onto a glass surface. B) *Plasmodium vivax* parasite development significantly decreased in female mosquitoes exposed to ATQ for 6 min. Prevalence (Prev.): two-sided chi-squared test, *n =* 249, degrees of freedom (df) = 1, χ2 = 133.3, \*\*\*\**P* < 0.0001. Intensity: two-sided Mann-Whitney *U* test, *n* = 249, df = 1, *U =* 706.5, \*\*\*\**P* < 0.0001. C) To evaluate the effect on sporozoites in the salivary glands: *Plasmodium vivax* parasite development significantly decreased the number of oocysts in female mosquitoes exposed to ATQ for 6min. Prevalence (Prev.): two-sided chi-squared test, *n* = 79 degrees of freedom (df) = 1, χ2 = 63.96, \*\*\*\**P* < 0.0001. Intensity: two-sided Mann-Whitney *U* test, *n* = 79, df = 1, *U =* 24, \*\*\*\**P* < 0.0001. The exposure method is shown in the graphic: orange represent ATQ coated onto a glass surface. D) *Plasmodium vivax* parasite development significantly decreased the number of sporozoites in female mosquitoes exposed to ATQ for 6 min. Prevalence (Prev.): two-sided chi-squared test, *n =* 91, degrees of freedom (df) = 1, χ2 = 57.58, \*\*\*\**P* < 0.0001. Intensity: two-sided Mann-Whitney *U* test, *n* = 91, *U* = 0, \*\*\*\**P* < 0.0001. Medians are indicated.

In additional experiments, tarsal exposure to ATQ for 6 min before infection again impaired oocyst survival (Fig 1C) and also significantly reduced sporozoites intensity and prevalence (Fig 1D).

Parasite prevalence and intensity of *P. vivax* infection were also significantly reduced when mosquitoes were exposed to ATQ 24h before (Fig 2A) or 12h after infection (Fig 2B). These findings indicate that ATQ can suppress *P. vivax* development in the female mosquitoes both before and after an infected blood meal.

**Fig 2.**
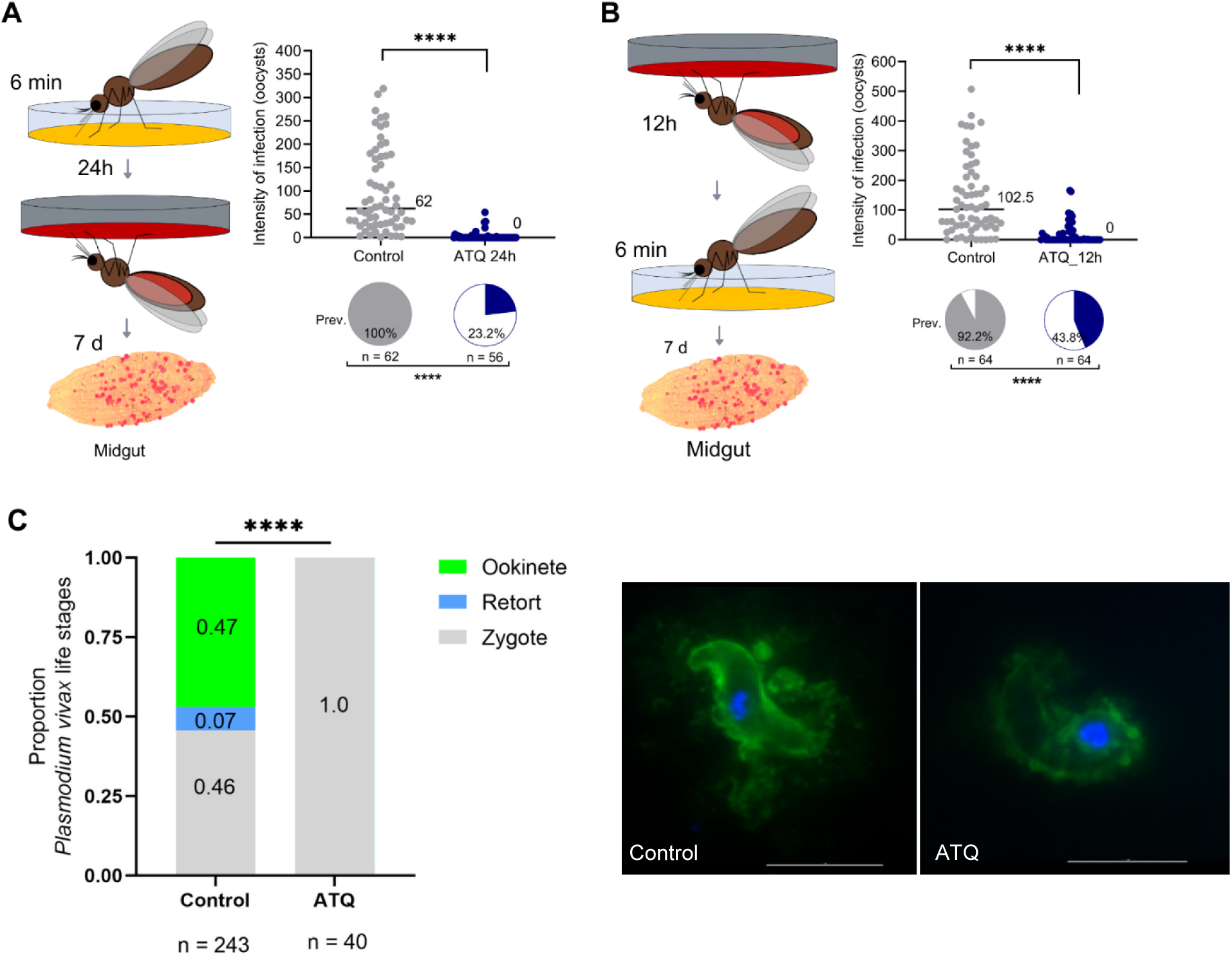
Atovaquone (ATQ) reduced *Plasmodium vivax* infection when mosquitoes were exposed for six minutes, either before or after infection. A) *Plasmodium vivax* prevalence and oocyst intensity were significantly reduced when female mosquitoes are exposed to ATQ (1 mmol per m^2^ for 6 min) 24 h before infection (prevalence: two-sided chi-squared test, *n =* 118, df = 1, χ^2^ = 74.90, \*\*\*\**P* < 0.0001; oocyst intensity: two-sided Mann-Whitney *U* test, *n* = 118, df = 1, *U =* 103.5, \*\*\*\**P* < 0.0001. B) Similar, prevalence and oocyst intensity were reduced when mosquitoes were exposed to ATQ 12 h after an infectious blood meal (prevalence: two-sided chi-squared test, *n =* 128, df = 1, χ^2^ = 34.49, \*\*\*\**P* < 0.0001; oocyst intensity: two-sided Mann-Whitney *U* test, *n* = 128, df = 1, *U = 562.5*, \*\*\*\**P* < 0.0001. Medians are indicated. C) Immunofluorescent assay of mosquito midgut lumens 21 h after *Plasmodium vivax* infection, using parasite-specific antibodies (anti-Pv25, green) and DNA staining (Hoechst, blue). Parasite forms observed in control group included mature ookinete and zygote (left), while ATQ-treated group displayed immature retort forms of ookinetes (right). Ten midguts were analyzed for each group. No retort forms and ookinetes were observed in ATQ-treated group, which exhibited only zygote (100% of parasites). In contrast, control group displayed a significant proportion of normal ookinetes (47%) and zygotes (46%), with retort forms constituting only 7% of the total parasites (Chi-square test (*n =* 243 number of parasites found at control and *n =* 40 number of parasites found at ATQ-treated, df = 2. χ^2^ = 40.72, *\*\*\*P* < 0.0001). Scale bar. 10 μm. As in Paton et al. [13], we confirmed that *P. vivax* parasites were killed at the early zygote-ookinete transition, as determined by immunofluorescence assay of infected midguts (Fig 2C). Additionally, ATQ-treated female mosquitoes had fewer parasites in the blood bolus when compared to controls (Fig 2C).

### No effect on *Plasmodium vivax* development in mosquitoes exposed to antimalarials used in Brazil

Considering the positive results of the direct contact assay using the antimalarial ATQ, we also tested antimalarials used in Brazil against *P. vivax*, such as primaquine (PQ), tafenoquine (TQ), chloroquine (CQ) and mefloquine (MQ) [21]. These were evaluated at the maximum exposure time of 60 minutes and at the same concentration as ATQ. The antimalarials PQ, TQ and CQ and the compound did not achieve blocking or reduction of infection (Fig 3A, B and C). Mefloquine-exposed mosquitoes had a significant reduction in oocyst intensity compared to the control (*P* = 0.0002), but the prevalence of infection was not affected (Fig 3D).

**Fig 3.**
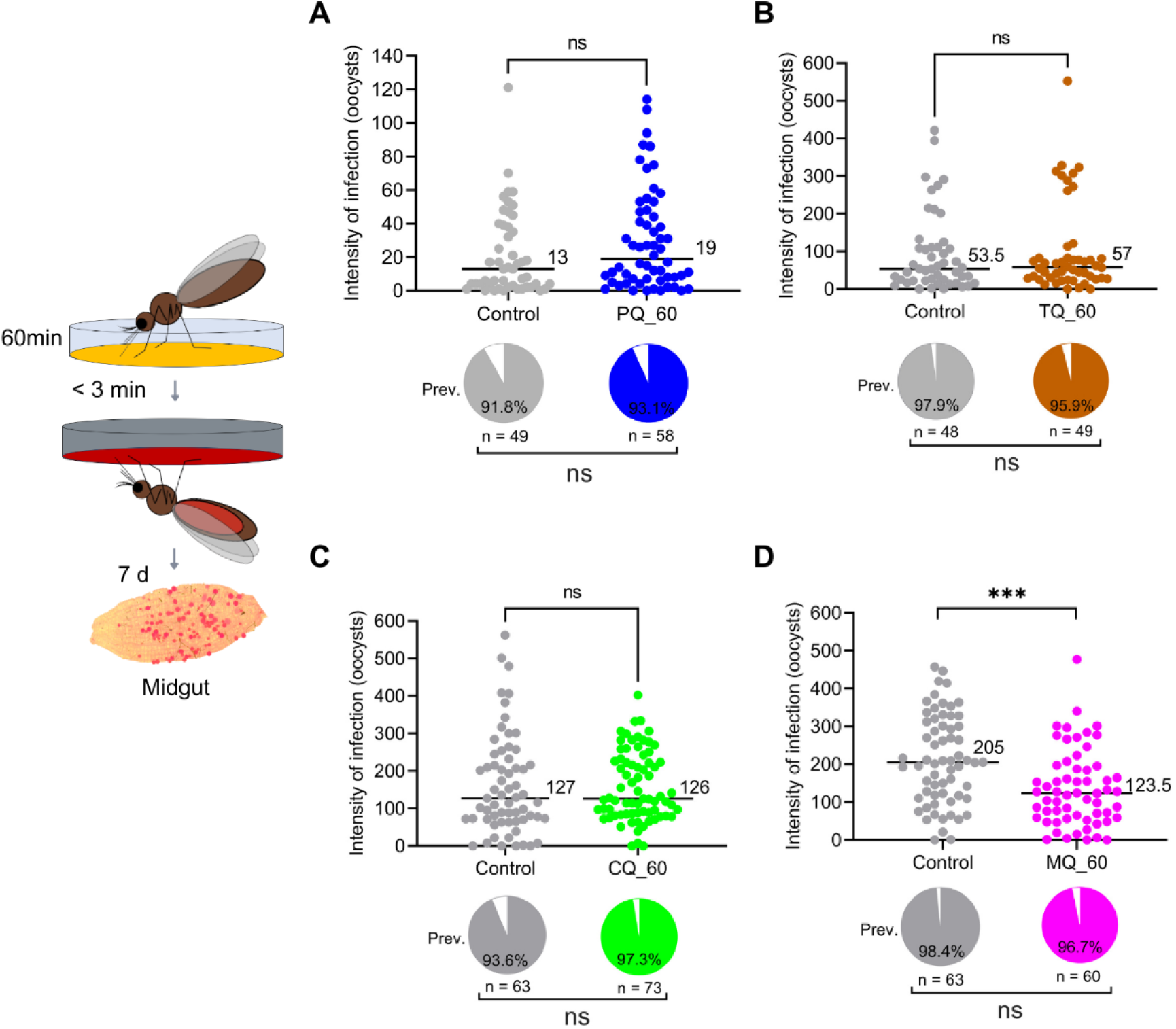
*Plasmodium vivax* antimalarials did not significantly affect parasite development in *Anopheles darlingi* mosquitoes were exposed to the following antimalarials A) Primaquine (PQ): *n* = 107, prevalence 93.1%, degrees of freedom (df) = 1, χ2 = 0.06161, *P* = 0.8040, ns. Oocyst intensity *P* = 0.1009, *U* = 1159, ns; B) Tafenoquine (TQ): *n* = 97, prevalence 95.9%, degrees of freedom (df) = 1, χ2 = 0.3231, *P* = 0.5698, ns. Oocyst intensity *P* = 0.7047, *U* = 1123, ns; C) Chloroquine (CQ): *n* =136, prevalence 97.3 %, degrees of freedom (df) = 1,, χ2 = 1.045, *P* = 0.3067; Oocyst intensity *P* = 0.4965, *U* = 2143, ns; except when exposed to D) Mefloquine (MQ): *n* = 123, prevalence 96.7%, degrees of freedom = 1, χ2 = 0.3937, *P* = 0.5303, ns; Oocyst intensity *P* = 0.0002, *U* = 1161, significant. Medians are indicated for all tests.

To investigate whether the physical barrier formed by the mosquito cuticle hindered the uptake of MQ after tarsal exposure, the drug was directly added to the infected blood meal at a final concentration of 10 μM and then offered to *An. darlingi* females. Atovaquone was used as a positive control. When added to the infected blood prior to infection, blocked/reduced oocysts development at the 7^th^ day post-DMFA (Fig 4A), and sporozoite infection at the 14^th^ day post-DMFA (Fig 4B). Mefloquine significantly reduced the oocyst and sporozoite intensity of infection (Fig 4C, D), showing that MQ retains partial transmission as in the tarsal exposition (Fig 3D) – highlighting its potential, albeit potent than ATQ, as a candidate for mosquito-target transmission-blocking strategies.

**Fig 4.**
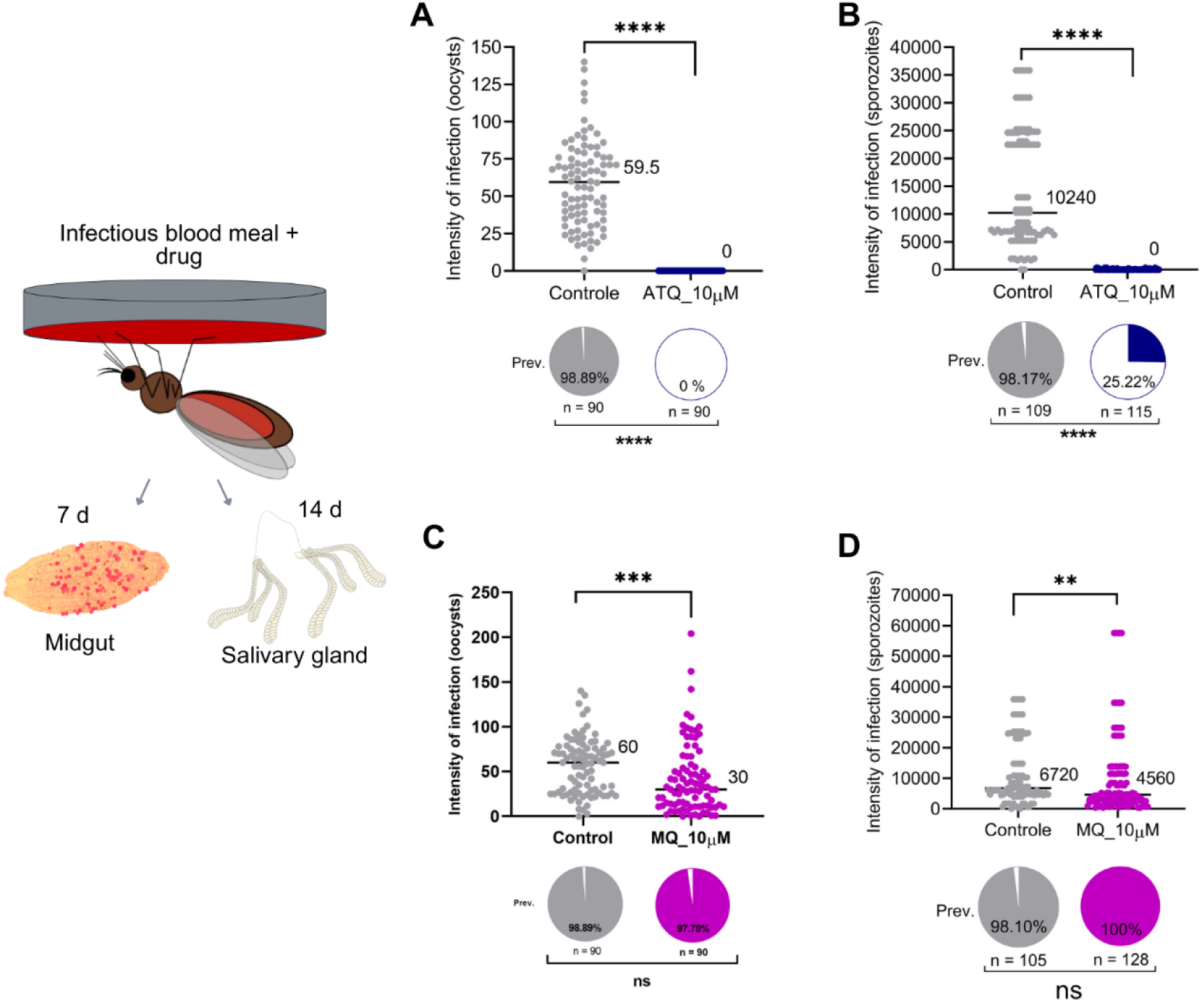
Antimalarials ATQ and MQ were added to blood infected with *Plasmodium vivax* and offered to female *Anopheles darlingi* mosquitoes via the Direct Membrane Feeding Assays (DMFA). A) Atovaquone (ATQ) added to the blood meal completely blocked *Plasmodium vivax* oocysts in the midguts and sporozoites in the salivary glands (B) of female mosquitoes. Oocyst prevalence: two-sided chi-squared test, *n* = 180, degrees of freedom (df) = 1, χ2 = 176.0, \*\*\*\**P* < 0.0001. Intensity: two-sided Mann-Whitney U test, *U* = 45, \*\*\*\**P* < 0.0001. B) Sporozoite prevalence: two-sided chi-squared test, *n* = 224, degrees of freedom (df) = 1, χ2 = 124.8, \*\*\*\**P* < 0.0001. Intensity: two-sided Mann-Whitney U test, *U* = 134, \*\*\*\**P* < 0.0001. C) The presence of Mefloquine (MQ) in the *Plasmodium vivax* blood meal partially reduced oocysts in the midguts of female mosquitoes and sporozoites in the salivary glands (D). Oocyst prevalence (C): two-sided chi-squared test, *n* = 180, degrees of freedom (df) = 1, χ2 = 0.3390, *P* = 0.5604. Intensity: two-sided Mann-Whitney U test, *U* = 2803, \*\*\**P* = 0.0003. D) Sporozoite prevalence: two-sided chi-squared test, *n* = 233, degrees of freedom (df) = 1, χ2 = 2.459 *P* = 0.1168. Intensity: two-sided Mann-Whitney U test, *U* = 5097, \*\**P* = 0.0015, Medians are indicated.

### Tarsal exposure of *Anopheles darlingi* to Nanchangmycin (NCG) does not reduce infection with *Plasmodium vivax*

Nanchangmycin (NCG) is a polyketide antibiotic [22] and was previously shown to block *P. vivax* development in *An. darlingi* when added to the infected blood prior to mosquito feeding [23]. Tarsal exposure of NGC to mosquito however did not significantly reduce the transmission of *P. vivax* to the vector (Fig 5).

**Fig 5.**
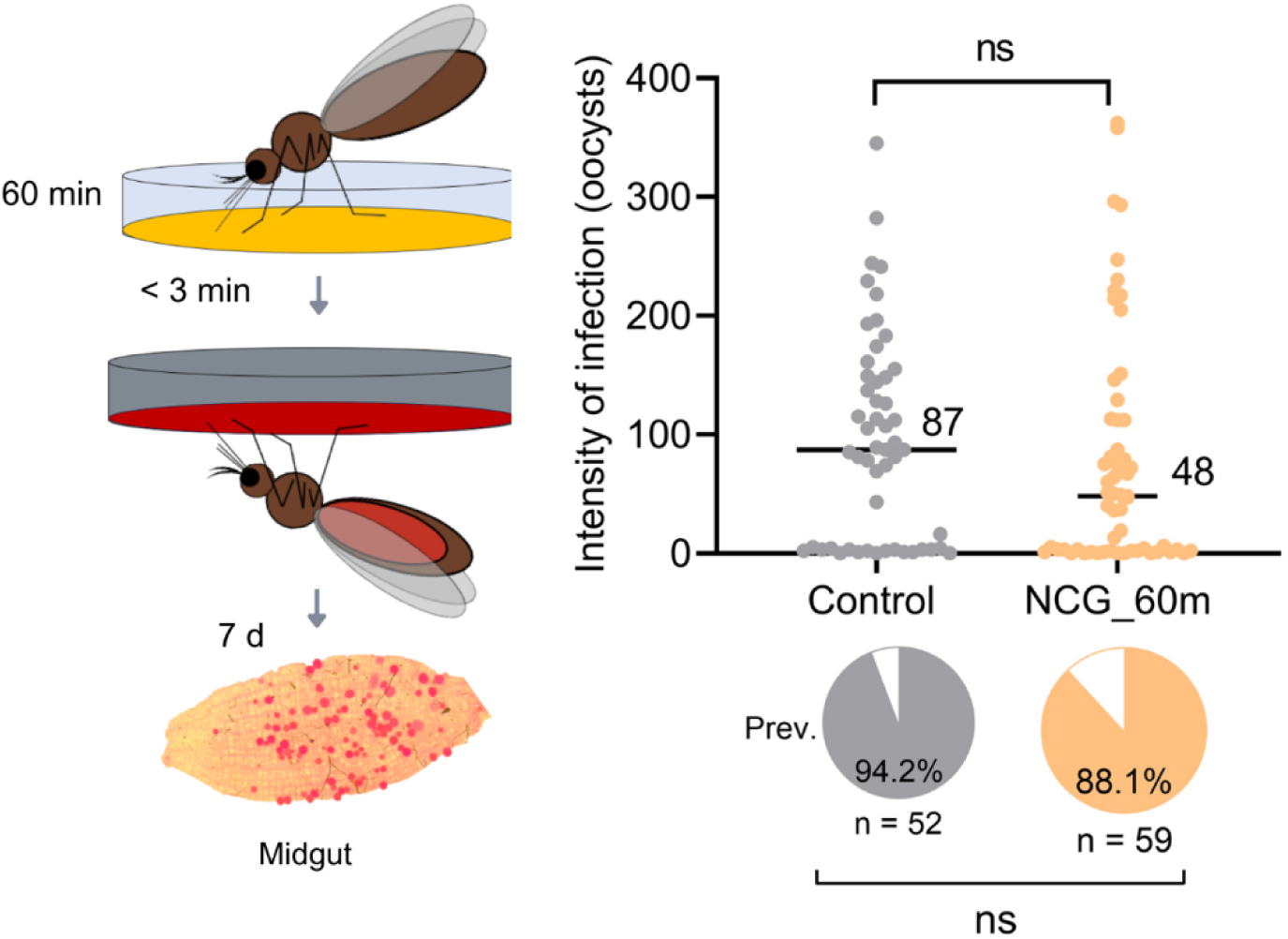
Tarsal exposure using Nanchangmycin (NCG) compound (1mmol per m^2^ for 60 min). *n* = 111, prevalence 88.14%. Prevalence (Prev.): two-sided chi-squared test, *n* = 111, degrees of freedom (df) = 1, χ^2^ = 1.253, *P* = 0.2630, ns. Intensity: two-sided Mann-Whitney *U* test, df = 1, *U* = 1249, *P* = 0.0916, ns. Medians are indicated.

### Antimalarials and Nanchangmycin (NCG) do not affect the survival of *Anopheles darlingi* females after tarsal exposure

Since some compounds, in addition to affecting *Plasmodium* development, may also impact the survival or overall fitness of mosquitoes. Therefore, we also assessed the survival of mosquitoes following tarsal exposure. The results indicated that mosquito survival on the 7^th^ day post-DMFA was not affected in any of the experimental groups exposed to tarsal treatment (Table 1).

**Table 1.** Survival of *Anopheles darlingi* submitted to tarsal exposure and infected with *Plasmodium vivax* on the 7^th^ day post-DMFA.

| Groups | Survival % (7<sup>th</sup> day post-DMFA) | P-value (Chi square) |
| --- | --- | --- |
| Control | 88.89 | 0.7649 |
| ATQ_60m | 90.36 |  |
| Control | 95.59 | 0.3921 |
| ATQ_6m | 97.54 |  |
| Control | 95.38 | 0.7258 |
| ATQ_6m_(24h.a.i.) | 96.67 |  |
| Control | 98.46 | 0.1167 |
| ATQ_6m_(12h.d.i.) | 92.96 |  |
| Control | 95.45 | 0.4524 |
| CQ | 92.40 |  |
| Control | 87.50 | 0.7283 |
| MQ | 89.55 |  |
| Control | 88.37 | 0.8018 |
| PQ | 89.70 |  |
| Control | 88.33 | 0.438 |
| TQ | 92.59 |  |
| Control | 94.91 | 0.5036 |
| NCG | 91.67 |  |

## DISCUSSION

Paton et al. [13] garnered significant attention within the scientific community with a landmark study demonstrating that incorporating ATQ into a glass substrate - on which blood-fed *Anopheles* mosquitoes rested - effectively eliminated *P. falciparum* parasites within the mosquitoes’ midgut blood meal. This novel approach, which delivers an antimalarial compound through surface contact during the mosquito’s resting phase pre- or post-blood-feeding, represents a highly innovative strategy to disrupt the *Plasmodium* transmission cycle and offers numerous advantages [24]. However, the effect of ATQ on the sporogonic development of *P. vivax* had yet to be evaluated. Here, we observed that the *P. vivax* development in *An. darlingi* mosquitoes was significantly impaired when mosquitoes were exposed to ATQ before or shortly after infection.

Numerous chemical compounds, including ATQ, have demonstrated efficacy against *P. falciparum* parasites, as well as against *P. berghei* and *P. yoelii* during sporogony [16,20,25–27]. However, data on the effects of ATQ on the sporogony development of *P. vivax* remain limited, as do studies on other antimalarials [28,29]. Atovaquone is well-known for its dual activity against both the initial liver and the pathogenic erythrocytic stages of *P. falciparum* and *P. vivax* [30,31]. Our *in vitro* experiments have shown that ATQ effectively eliminates asexual stages (S2 Table) and ookinetes (S3 Table) of *P. vivax* clinical isolates obtained from patients. Due to its strong activity, ATQ has been used as a positive control in some of our *in vitro* assays.

Notably, ATQ was the only antimalarials among those tested that was capable of penetrating the cuticle of mosquito legs via tarsal contact, directedly reaching the hemolymph and blocking *P. vivax* transmission in *An. darlingi*. For this strategy to work, compounds/antimalarials must overcome the exoskeleton barrier to access internal tissues where the parasite develops. To achieve this, the compounds should possess specific characteristics [15]. First, it must be lipophilic; with a positive log*P* value, as lipophilicity is a critical factor in a compound’s absorption, distribution, membrane penetration, and overall pharmacokinetic properties (ADME: absorption, distribution, metabolism, and excretion). Lipophilicity is a key parameter used in pharmaceutical and biotech industries to evaluate drug efficacy. Second, the polar surface area (PSA) of the compound should not be excessively large, as larger molecules may have to overcome cuticle penetration resistance [13]. Moreover, the compound must exhibit intrinsic antiplasmodial activity.

Atovaquone, a ubiquinone analog, inhibits the mitochondrial electron transport chain by displacing ubiquinone, thereby disrupting ATP synthesis and *de novo* pyrimidine biosynthesis, ultimately leading to parasite growth inhibition [32]. Given the effectiveness of ATQ, future studies will evaluate a dose-response effect and analytically determine time kinetics of ATQ dissemination through the cuticle and persistence in the midgut and salivary glands. Furthermore, ATQ is highly lipophilic, with a positive log*P* and a PSA of 54.4 Â^2^ (S4 Table). Based on immunofluorescence assays using anti-Pvs25 [33], our results suggest that ATQ also targets *P. vivax* during the early zygote-ookinete transition, as previously shown for *P. falciparum* [13]. This is consistent with the parasite’s development window, 18-24 hours post-infection, when ookinetes are typically formed in the midgut (Fig 2C).

To further explore ATQ’s mechanism, we investigated its interaction with the cytb target using molecular docking. Although an experimental structure of *P. vivax* cytb is not available, the Alphafold-predicted model (AF-O63696-F1-v4), shows high structural similarity to *Saccharomyces cerevisiae* cytbc1 (PDB ID: 4pd4) which contains ATQ co-crystallized in Qo binding site (S1 Fig). Structural alignment between the two proteins results in a Root Mean Square Deviation (RMSD) of 0.824 Â, indicating strong conservation, particularly within the binding pocket. This justified the use of the *S. cerevisiae* structure for docking validation via redocking, followed by docking simulations on the *P. vivax* model using the same parameters (S1 Fig). The docking results in *P. vivax* cytb revealed ATQ binding to a conserved hydrophobic pocket involving key residues such as Phe123, Met133, Trp136, Gly137, Val140, Ile258, Leu285, Leu288, Pro260, Phe264, Tyr268, Leu271, Ile141, Phe267, and Val284) (S2 Fig), according to the residues described by Birth et al. [34], in which the position of the residues between cytb of *P. vivax* and chain C of cytbc1 of *S. cerevisiae* differ due to the alignment performed on the sequences in addition to four residues that also differ between the proteins (S3 Fig). The influence of the differences of these four residues on the ATQ binding site will be investigated in future work through molecular dynamics simulations. The binding energy was – 9.825 kcal/mol in *P. vivax* model, confirming a favorable interaction. In addition, electrostatic surface analysis supported the physicochemical compatibility of ATQ with the binding site in *P. vivax*. The strong structural and electrostatic conservation of the Qo binding site between *S. cerevisiae* structural and *P. vivax* reinforces the validity of this comparative modeling approach and supports the use of the *P. vivax* cytb model for future *in silico* screening of transmission-blocking compounds.

Mefloquine, another antimalarial tested, affected oocyst and sporozoite intensity but not infection prevalence (Fig 4C and D). Commonly used for prophylaxis and combination therapies [35], MQ is a 4-methanolquinoline structurally related to quinine [35,36]. Its proposed mechanism involves inhibition of heme detoxification [37–39]. Interestingly, a study on the inhibition of esophageal carcinoma cell growth *in vitro* observed the downregulation of protein expression in all subunits involved in oxidative phosphorylation. Proteomic analysis indicated that mitochondria are particularly affected by MQ [40], including the bc1 complex, pointing to similarities to the mode of action of ATQ. A previous study conducted by Li et al. [41] also highlighted MQ’s ability to inhibit mitochondrial respiration. Mefloquine exhibits stage-specificity action similar to quinine, primarily targeting large ring and trophozoite stage asexual parasites [35,37,39]. There is also some evidence for sporontocidal activity. For instance, Coleman et al. [42] showed dose-dependent effects on *P. berghei* ANKA sporogony in *An. stephensi*, with changes in oocyst numbers and the extent of sporozoite invasion into salivary glands. Further research is needed to fully elucidate the mechanisms underlying MQ’s effects on parasite development. Mefloquine’s physicochemical properties permit cuticle penetration (S4 Table), unlike TQ, PQ and NCG, all of which have PSA> 60 Â^2^.

Interestingly, NCG completely blocked transmission of *P. vivax* in *An. darlingi* and *P. falciparum* in *An. stephensi* during DMFA and SMFA assays, respectively [23], but had no effect when administered to mosquitoes via tarsal exposure likely due to its higher PSA (S4 Table). Alternative delivery methods, such as sugar baits (ATSBs), could be explored such compounds [20]. Nanchangmycin is a polyether ionophore antibiotic produced by *Streptomyces nanchangensis* [22] and has known insecticidal properties against silkworms and anti-bacterial activity *in vitro* [43,44].

Chloroquine, although possessing a low PSA (S4 Table), which likely allows it to penetrate the cuticle of mosquito legs via tarsal contact and reach the hemolymph, has not showed evidence of sporontocidal activity. Measuring compound concentrations in mosquito hemolymph using HPLC after tarsal exposure could clarify pharmacokinetics profile and drug-*Plasmodium* interactions in mosquitoes [45,46]. Paton et al. [13] demonstrated that other cytochrome *b* inhibitors like decoquinate (DEC) and pyrimethamine (PYR) were ineffective in blocking *Plasmodium* development in exposed mosquitoes, likely due to higher PSA. Recently, harmane – a small, hydrophobic β-carboline secreted by *Delfitia tsuruhatensis* - was found to fully inhibit *P. falciparum* development via tarsal penetration [47]. Harmane has small PSA (28,7 Â^2^; PubChem, 2024), which likely supports this trans-cuticle uptake.

Regarding potential mosquito fitness costs, Paton et al. [13] reported no effect in survival rate and fecundity of *Anopheles* mosquitoes at 48 hours after 60 minutes of exposure to concentrations up to 1 mmol/m^2^ via tarsal contact. In our experiments, mosquito survival was evaluated until the 7^th^ day post-contact and *P. vivax* infection, and no reductions in survival were observed for any compounds tested. These results suggest that ATQ, and even MQ, selectively target *P. vivax* without compromising mosquito viability.

Despite the promising effects of ATQ and MQ on *P. vivax* development in *An. darlingi*, it is crucial to recognize the limitations of applying human-use antimalarials for vector-targeted control. Malaria parasites have evolved resistance to nearly all antimalarial used in humans. Therefore, it would be naïve to assume that compounds targeting parasites during sporogony will not face resistance evolution. Kamiya et al. [20] proposed that using different compounds in humans and mosquitoes could reduce selective pressure for resistance and drug combination was shown to be highly effective against mosquito stages of *P. falciparum* in a recent study [15]. Our proof-of-concept study showing that with ATQ blocking *P. vivax* in *An. darlingi* provides a foundation for identifying compounds with structural and chemical similarities to ATQ for further evaluation. Using dedicated compounds to target parasites during sporogony offers an additional control strategy, which could reduce the reliance on human antimalarials for suppressing transmission. This approach represents a promising avenue for integrated malaria control, combining interventions in both humans and mosquitoes to achieve sustainable transmission reduction.

## Materials and methods

### *Anopheles darlingi* Direct Membrane-Feeding Assays (DMFA)

Females *An. darlingi* mosquitoes were obtained from the colony established at the Malaria Vectors Production and Infection Platform (PIVEM), located at FIOCRUZ-RO, Brazil, as described by Araujo et al. [48]. Mosquitoes were raised at a temperature of 26°C ± 1°C and a relative humidity of 70 ± 10%, being fed a 15% honey solution.

For the experiments, 25-50 female *An. darlingi* mosquitoes, 3-5 days old, were used for each experimental and control group, with sucrose food removed the day before the DMFA.

### Tarsal exposure test

For tarsal exposure, the compound or antimalarial were diluted in an appropriate volatile vehicle. Solutions were prepared considering the area of the Petri dish and the molecular weight of the compound or antimalarial (S4 Table) [13].

One milliliter of the prepared solution at a concentration of 1 mmol per *m*^2^ (*mg*/*m*^2^) was pipetted onto the Petri dish. The volatile vehicle used for compound dilution was used as control. The plates were then kept overnight on an orbital shaker at 25°C and 100 rpm to coat the entire area of the plate. A transparent plastic container of the same diameter as the Petri dish was chosen to allow the two parts to fit together. This container was then adapted to introduce the mosquitoes, ensuring that their tarsi were in contact with the compound or antimalarial-impregnated surface of the plates. These procedures were performed as described by Paton et al. [13].

The antimalarials tested in this tarsal exposure assay were atovaquone (ATQ), primaquine (PQ), chloroquine (CQ), mefloquine (MQ), tafenoquine (TQ), and the compound nanchangmycin (NCG), with an initial exposure time of 60 minutes before infection (60 m.b.i.). Direct membrane feeding assay was then performed as described below.

When the exposure time of 60 min was efficient to block mosquito infection, the time was reduced to 6 minutes. It also was tested mosquito exposure to the compounds 24 h before or 12 h after infection.

### Blood collection from patients with diagnosed *Plasmodium vivax* infection

Individuals who participated in the study were selected from patients diagnosed with vivax malaria through Giemsa-stained blood smears collected at the Center for Tropical Medicine Research (CEPEM) in Porto Velho, Rondônia, an endemic region in the Brazilian Amazon (under protocol approval number #28176720.9.0000.0011). Volunteers were selected based on the following criteria: monoinfection with *P. vivax* by thick blood smear (parasitemia > 2000 parasites/ µL), age between 18 and 85 years, without signs or symptoms of severe malaria or concomitant diseases, with or without a previous history of malaria, no pregnant, and agreed to the study procedures.

### Direct Membrane-Feeding Assays

Patient’s venous blood was collected in lithium heparin tubes (10 mL, Vacutainer, BD) and then transported in a thermal bottle at 37°C from CEPEM to PIVEM. The heparinized tube was centrifuged at 1,500 rpm for 10 minutes. Only the red blood cells were used. Prior to mosquito feeding, 500 µL of inactive AB^+^ serum was mixed with 500 µL of parasitized red blood cells obtained [49].

The prepared blood was offered to the female mosquitoes of each group for 30 minutes using glass feeders attached to a water bath or disc with the Hemotek^®^ device, as previously described [48]. After this period, unfed or partially fed mosquitoes were removed, leaving only fully engorged mosquitoes in the experimental cages for subsequent examination of sporogonic development. A cotton wad soaked in a 15% honey solution was regularly provided and changed every two days until mosquito dissection. The mosquito survival was evaluated daily until the 7^th^ day after blood feeding, when the midgut was dissected. To determinate the sporozoite intensity of infection, salivary glands were dissected at the 14^th^ day post-DMFA.

For dissection, mosquitoes were anesthetized on ice, immersed in 70% ethanol, and transferred to phosphate-buffered saline (1X PBS). The midguts were stained with 0.2 % mercurochrome solution and examined for the presence of oocysts by microscopy. For experiments involving the addition of antimalarials to *P. vivax*-infected blood, which was then offered orally to female *An. darlingi*, the antimalarials were first mixed with the inactivated AB^+^ serum (5 μL of antimalarial + 495 μL of serum). Patient-derived red blood cells (separated by centrifugation, as previously described) were then admixed, resulting in final volume of 1,000 μL with 10μM of the antimalarial.

### Ookinete immunofluorescence assays – IFA

Twenty-one hours after an infectious blood meal, ten female mosquitoes from control and ATQ groups were aspirated into 1x PBS at 4° C. Midguts with blood bolus were isolated and transferred to 20 μL of 1X PBS on ice. Guts were disrupted by pipetting and the crude isolate homogenized by vortexing briefly (about 5 seconds), and 10 μL of the homogenate was spotted onto a poly-L-lysine-coated slide and air dried. Once dry, the tissues were fixed by incubation with 4% paraformaldehyde (PFA) for 10 minutes [13]. Slides were then rinsed three times with 5 to 10 μL of 1x PBS, to blocked with 1% BSA in 1 x PBS for 1 h and then rinsed again three times with 5 to 10 μL 1x PBS. Ookinetes were stained with mouse antibody raised against the *P. vivax* surface protein Pvs25 (100 μg/mL) for 1 h in a humid box, at room temperature [33]. Secondary staining was carried out with 1:100 dilution of Alexa Fluor 488 goat anti mouse IgG (Invitrogen) for 1 h a dark humid box. After several washes with 1 x PBS, the cells were counter stained with Hoechst 33342 (10 μg/mL), washed and then the tissues were mounted in Everbrite mounting medium (Biotium). Slides were examined by fluorescence microscopy (Nikon Eclipse 80i^®^) with 100x oil immersion objective and the images were captured using the *software* Nikon Nis Elements^®^.

### Statistics analysis

Data of infection intensity were analyzed using Mann-Whitney, while infection prevalence and the proportion of parasite life stages (zygote, retort and ookinete) were analyzed using the Chi-square test. Experiments in which the controls presented medians lower than 2.5 oocysts per mosquito and infection prevalence below 60% were not taken into consideration in the statistical analyses (S1A and S1B Table). The transmission-reducing assay (TRA) was measured as percentage of reduction in oocyst density and the transmission-blocking assay (TBA) was evaluated as inhibition in the prevalence of infected mosquitoes [48,50] (S1C Table).

Female mosquitoes exposed to antimalarials and the NCG compound were monitored until the seventh day after infection to verify survival with the Kaplan-Meier survival curve and the survival rate was compared using the Log-rank test.

All analyses were performed using *GraphPad Prism software* (version 9.3.1).

## Acknowledgments

We thank the staff of the Malaria Outpatient Clinic at the *Centro de Pesquisa de Medicina Tropical de Rondônia* (CEPEM) in Porto Velho/Rondônia/Brazil for recruiting study participants and collecting blood samples, as well as the volunteers who donated blood for this study.

## Supporting information captions

**S1 Table. Individual and summary data from Direct Membrane Feeding Assays (DMFA).** A) Transmission-reducing activity (TRA, %) and transmission blocking activity (TBA, %) for each DMFA. B) Individual DMFA results. C) Summary of DMFA data.

**S2 Table**. **Characteristics of *Plasmodium vivax* obtained from patients and used in *ex vivo* assay with Atovaquone (ATQ).**

**S3 Table. Characteristics of *Plasmodium* vivax obtained from patients and used in ookinete inhibition assay with Atovaquone (ATQ).**

**S4 Table**. **Chemical properties from all compound’s testes in this study – atovaquone, primaquine, tafenoquine, chloroquine, mefloquine and nanchangmycin.**

**S1 Fig. Structural alignment - Structural superposition of *Plasmodium vivax* cytochrome b (light pink, AlphaFold model AF-O63696-F1-v4) and the C-chain of *Saccharomyces cerevisiae* cytochrome bc1 complex (green, PDB ID: 4pd4). Atovaquone (ATQ) is shown in yellow sticks as positioned in the crystallographic structure of *Saccharomyces cerevisiae*. The Root Mean Square Deviation (RMSD) between the two models was 0.824 Å, indicating high structural similarity. Structural visualization was performed in ChimeraX.**

**S2 Fig. Docking result:** Atovaquone (ATQ) binding site on the cytochrome b (AlphaFold ID: AF-O63696-F1-v4). A) Tertiary structure of the cytochrome b (AlphaFold ID: AF-O63696-F1-v4) shown as a light pink ribbon. A transparent surface highlights the overall shape of the protein. ATQ is represented in yellow sticks. B) Close-up of ATQ-binding residues shown in ball-and-stick representation with residue names labeled (Phe123, Met133, Trp136, Gly137, Val140, Ile258, Leu285, Leu288, Pro260, Phe264, Tyr268, Leu271, Ile141, Phe267, and Val284). C) Electrostatic surface potential of the ATQ binding pocket. Red indicates negatively charged regions, blue indicates positively charged regions, and white indicates neutral areas. Structural visualization was performed in ChimeraX.

**S3 Fig. Sequence alignment:** Sequence alignment of *Plasmodium vivax* cytb proteins (AF-O63696) and *Saccharomyces cerevisiae* Cytb C chain (PDB ID: 4pd4), performed using Clustalw. Residues highlighted in yellow correspond to those involved in atovaquone (ATQ) binding. Green arrows indicate residues that changed between proteins. Image made in ESPript 3.0.

